# Epithelial-Mesenchymal Plasticity is Associated with Immunosuppressive Features in Canine Mammary Carcinomas

**DOI:** 10.1101/2025.09.19.677414

**Authors:** Kimaya Bakhle, Sophie Nelissen, Lynna Li, Ioannis Ntekas, Iwijn De Vlaminck, Santiago Peralta, Andrew D. Miller, Anushka Dongre

**Affiliations:** Department of Biomedical Sciences, College of Veterinary Medicine, Cornell University, Ithaca, NY 14853, USA; Meinig School of Biomedical Engineering, Cornell University, Ithaca, NY 14853, USA; Department of Clinical Sciences, College of Veterinary Medicine, Cornell University, Ithaca, NY 14853, USA; Department of Population Medicine and Diagnostic Sciences, College of Veterinary Medicine, Cornell University, Ithaca, NY 14853, USA

**Keywords:** epithelial-mesenchymal plasticity, tumor microenvironment, comparative oncology, canine mammary carcinoma, breast cancer

## Abstract

Epithelial-mesenchymal plasticity (EMP) is a cellular process activated in carcinoma cells to drive invasiveness, metastasis, and chemoresistance. Recently, we have demonstrated that activation of the EMP program also results in the assembly of an immunosuppressive tumor microenvironment and resistance to immunotherapy in an immunocompetent orthotopic murine model. However, it has yet to be shown whether these findings can be translated to canine carcinomas. Here, we show that in spontaneous canine mammary carcinomas (CMCs), which share similar histopathologic, molecular, and clinical features with human breast cancers, EMP activation is linked to the recruitment of immunosuppressive cells including regulatory T-cells and M2-like macrophages. Additionally, through transcriptomic profiling of CMCs, we identify that the glycoprotein CD109 is associated with EMP-mediated immunosuppression across canine, murine, and human breast cancer models. CD109 has been previously associated with tumorigenicity, but not immunosuppression in cancers of any species. Finally, we identified upregulation of several immunosuppressive paracrine factors across multiple canine carcinomas, including oral squamous cell carcinoma, urothelial carcinoma, and pulmonary carcinoma samples. These findings demonstrate for the first time that the EMP program is associated with immunosuppression in canine carcinomas, with direct translational implications for human breast cancers.

## Introduction

The epithelial-mesenchymal transition (EMT) is a cell-biological process that enables epithelial cells, which express the markers EpCAM and E-cadherin, to gain mesenchymal markers, such as vimentin and N-cadherin, and features including invasiveness, metastatic potential, and stem-cell like properties (1–8). In neoplasia, these malignant features lead to recurrent, invasive primary tumors as well as notoriously incurable metastatic disease (6, 9–11). These changes are driven by EMT-inducing transcription factors (EMT-TFs) Zeb1, Twist, Slug, and Snail. EMT-TFs are activated by several signaling molecules secreted by stromal and immune cells within the tumor microenvironment (TME) (12–16). These molecules include TGF-β, WNT ligands, NOTCH receptor ligands, epidermal growth factor (EGF), fibroblast growth factor (FGF), hepatocyte growth factor (HGF), platelet-derived growth factor (PDGF), IL-6, CCL18, and tumor necrosis factor (TNF) (8). Importantly, the EMT program is a dynamic process that results in a spectrum of hybrid or quasi-mesenchymal states, which express both epithelial and mesenchymal markers. Moreover, mesenchymal cells can revert to the epithelial state by activating the reverse process of mesenchymal-to-epithelial transition. Accordingly, this program is increasingly being recognized as epithelial-mesenchymal plasticity (EMP) (8, 17–20).

We have previously identified that EMP within murine breast cancers determines responses to immune attack (21, 22). We established novel, preclinical murine models of epithelial and quasi-mesenchymal mammary tumors, using cell lines derived from the autochthonous mouse mammary tumor virus-polyoma middle T antigen (MMTV-PyMT) model. Using this model, we determined that epithelial tumors assembled an immunopermissive TME, in which the tumors were densely infiltrated by CD8^+^ T-cells and anti-tumor M1-like macrophages were restricted to the tumor periphery (8, 21). Conversely, quasi-mesenchymal tumors excluded exhausted CD8^+^ T-cells to the periphery and instead recruited regulatory T-cells (Tregs) and pro-tumor M2-like macrophages (21). Findings of EMP-associated immunosuppression have also been reported in murine models of other cancer types, including melanoma, lung adenocarcinoma, and hepatocellular carcinoma (23– 25). These alterations in the TME also resulted in differences in immunotherapy response, such that epithelial mammary tumors were responsive to anti-CTLA-4 immune checkpoint blockade (ICB) therapy, but quasi-mesenchymal mammary tumors were resistant to this same treatment. Furthermore, mixed epithelial-mesenchymal tumors containing a minority fraction of quasi-mesenchymal cells (10%) were also almost completely resistant to anti-CTLA-4 therapy (21). This is of great clinical importance, as heterogeneous tumors in breast cancer patients may contain minority fractions of quasi-mesenchymal cells that could drive resistance of these tumors to multiple forms of ICB therapy (26–30).

To identify drivers of EMP-mediated immunosuppression, we previously performed a transcriptomic analysis of paracrine factors expressed by epithelial and quasi-mesenchymal murine mammary cancer cell lines. We identified genes that were upregulated in two independent murine quasi-mesenchymal cell lines compared to their epithelial counterparts and associated with paracrine immunosuppressive signaling (hereafter referred to as EMP-related paracrine factors) (21, 22) This study identified that in comparison to epithelial cells, the quasi-mesenchymal cancer cells upregulated several immunosuppressive factors including SPP1, MASP-1, CXCL12, CSF1, CD73, LGALS3, and TGF-β1 (22). Of these, abrogation of the macrophage chemoattractant CSF1 or the cytokine SPP1 from quasi-mesenchymal cancer cells partially sensitized the tumors to anti-CTLA-4 immune checkpoint blockade therapy. Moreover, knockout of CD73 from quasi-mesenchymal cancer cells completely sensitized these tumors to anti-CTLA-4 therapy (22). This was the first report of completely eliminating quasi-mesenchymal cancer cells by targeting a single factor (22). This potent immunosuppressive role can be attributed to the generation of adenosine by CD73. Adenosine acts through a variety of receptors to promote pro-tumor immune cell functions (31–35). This is clinically significant, as quasi-mesenchymal cancer cells lead to recurrent, metastatic disease (8). These studies have elucidated an important mechanism of immunosuppression and immunotherapy resistance in breast tumors, however it is yet to be shown whether these patterns are conserved in other species. Determining whether EMP induces expression of immunosuppressive factors in naturally occurring tumors from other species would provide rationale for targeting these factors for therapeutic benefit.

In this study, we employed naturally occurring canine tumors to overcome limitations posed by murine models. Specifically, rodent models of cancer lack key features of spontaneous disease, do not fully recapitulate the heterogeneity of human cancers, and are housed under specific-pathogen free conditions, which markedly impacts immune development. Naturally occurring canine cancers are more representative of disease progression in humans (36–39). Dogs are also immunologically experienced, with exposure to vaccinations and infections before the development of cancers. The duration of the immune response to spontaneous cancer in canines is much longer compared to rodents, and therefore better recapitulates the development of antitumor immunity in humans (39). Cancers in dogs are treated in the same ways as humans: surgery, radiation therapy, and chemotherapy. Clinical trials in these companion animals can be conducted in a shorter time frame compared to humans, which can accelerate the progression of therapeutics from preclinical to human clinical trials (39). Logistically, accessing canine tumor samples involves far fewer hurdles compared to human tumor samples. Moreover, the associated medical records for these canine patients are often available to correlate clinical information and treatment response with the biospecimen. Similar to breast cancers in women, canine mammary carcinomas (CMCs) are one of the most common neoplasm in dogs and are associated with invasiveness, metastasis, and recurrence (40–43). CMCs share features with human breast cancer, including histologic subtypes, molecular subtypes, clinical features, and metabolic alterations, making them an excellent model for studying this disease (36, 44–49).

Here, we show that EMP status is associated with advanced histologic grade as well as an immunosuppressive TME in CMCs. Furthermore, transcriptomic profiling of CMCs across a spectrum of EMP states shows that heterogeneous and quasi-mesenchymal CMCs upregulate immunosuppressive factors, including the glycoprotein CD109. Finally, using RNA sequencing (RNA-seq) datasets from additional canine tumors of epithelial origin including oral squamous cell carcinoma, pulmonary carcinoma, and invasive urothelial carcinoma, we show conserved EMP-induced expression of immunosuppressive paracrine factors. Our findings suggest that EMP-driven immunosuppression is observed in several canine carcinomas, with translational relevance for humans. Additionally, we have identified EMP-associated factors that may be useful as biomarkers or therapeutic targets for detection and treatment of advanced, heterogeneous, or quasi-mesenchymal tumors in both canine and human patients.

## Results

### Canine mammary carcinomas of advanced histologic grade show increased epithelial-mesenchymal plasticity and immunosuppression

We first sought to identify if the EMP status of CMCs was associated with advanced histologic grade (I/III: low-grade, II/III: intermediate-grade, III: high-grade). We selected a total of 52 CMC samples of varying histologic grade and diagnosis **(Table S1)**. The samples came from client-owned female dogs with a median age of 10 years, and the majority of the dogs were spayed. Since the EMP program is associated with invasive and metastatic capabilities and CMC grade is assigned based on tubule formation, nuclear pleomorphism, and mitotic activity, we determined if tumors of higher grade would have decreased expression of the epithelial marker E-cadherin and increased expression of the mesenchymal marker vimentin, signifying activation of the EMP program in these tumors (48). Accordingly, we found that vimentin labeling increased with histopathological grade, both as a percentage of marker-expressing neoplastic cells and a composite score accounting for the percentage and intensity of staining **(Fig 1A-C, S1-2A)**. Furthermore, we found that neoplastic cells in grade I, vimentin-low tumors displayed a cuboidal to columnar cellular morphology with E-cadherin localizing to the membrane **(Fig 1A)**. Conversely, neoplastic cells in grade III, vimentin-high tumors displayed a partial (an intermediate state displaying a hybrid of epithelial and mesenchymal features) to spindle-shaped morphology, as well as cytoplasmic translocation of E-cadherin **(Fig 1A)**. Finally, the grade I tumors maintained some degree of tubular formation, whereas this structure was gradually lost in grade II and III tumors **(Fig 1A)**. Since myoepithelial cells and neoplastic epithelial cells undergoing EMP can both co-express E-cadherin and vimentin, we excluded 21 CMCs with overt myoepithelial differentiation. The remaining 31 CMCs showed even stronger associations between histopathological grade and upregulation of vimentin, as well as co-localization of E-cadherin and vimentin **(Fig 1B, S2B)**. These findings suggest that CMCs of increased grade activate the EMP program to promote malignant, invasive features.

**Figure 1.**
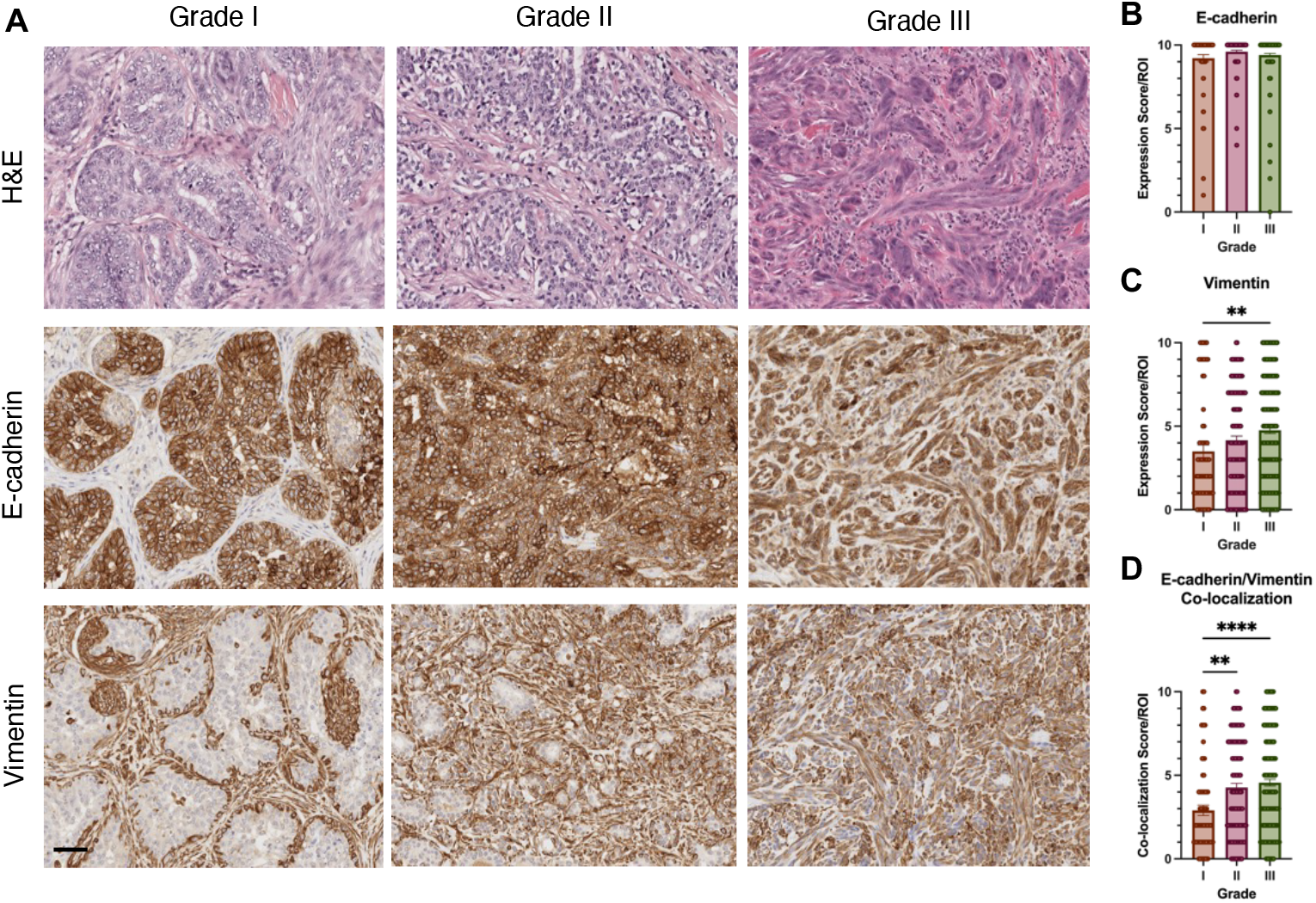
High-grade canine mammary carcinomas (CMCs) activate epithelial-mesenchymal plasticity (EMP). **(A)** Fifty-two archived formalin-fixed paraffin-embedded (FFPE) CMC specimens were acquired from the New York State Veterinary Diagnostic Laboratory. The samples were labeled for EMP markers E-cadherin and vimentin using immunohistochemistry. **(B-C)** Twenty-one CMC specimens with overt myoepithelial differentiation were excluded from this analysis. For the remaining 31 CMC specimens, percentage of neoplastic cells expressing E-cadherin and vimentin was scored on a 0-10 scale and across 15 high-power fields (HPF) for each tumor. **(D)** Estimated co-localization of E-cadherin and vimentin labeling in neoplastic cells of CMCs. All graphs are plotted as the mean value with error bars indicating the standard error of the mean. Data analyzed using one-way ANOVA: **** p < 0.0001, ** p < 0.01, ns (p > 0.05) if not indicated. Scale bar indicates 100 µm. ROI = region of interest.

Next, we aimed to determine whether these high grade, quasi-mesenchymal tumors assembled an immunosuppressive TME compared to the low grade, epithelial-like tumors. Based on previous findings from murine models, we hypothesized that EMP activation would convert the immunopermissive TME of epithelial tumors into an immunosuppressive TME in quasi-mesenchymal tumors (21, 23–25, 50). Indeed, we found that the number of tumor-infiltrating CD3^+^ T-cells decreased as histologic grade increased, with grade I, more epithelial tumors showing the highest infiltration and grade III, quasi-mesenchymal tumors showing the least infiltration **(Fig 2A-B, S3)**. This is clinically relevant because the presence of tumor-infiltrating T-cells directly correlates with response to immunotherapies in human patients (33, 51, 52). Although we did not see differences in the numbers of FOXP3^+^ Tregs, the ratio of FOXP3^+^ Tregs/CD3^+^ T-cells was significantly increased in both grade II and III tumors compared to grade I tumors **(Fig 2C-D, S3)**. Similarly, we found that although the number of Iba1^+^ total macrophages and CD204^+^ M2-like macrophages did not correlate with grade, the ratio of CD204^+^ M2-like macrophages/Iba1^+^ macrophages increased in grade III tumors compared to grade I tumors **(Fig 2E-G, S3)**. These findings suggest that more invasive CMCs of advanced histologic grade are associated with EMP activation, as well as recruitment of increased proportions of immunosuppressive M2-like macrophages and Tregs. This is consistent with the patterns that we, and others, have observed in murine models of epithelial and quasi-mesenchymal tumors (21–23, 25, 50, 53). Taken together, these findings demonstrate that EMP is associated with immunosuppression in canine mammary carcinomas.

**Figure 2.**
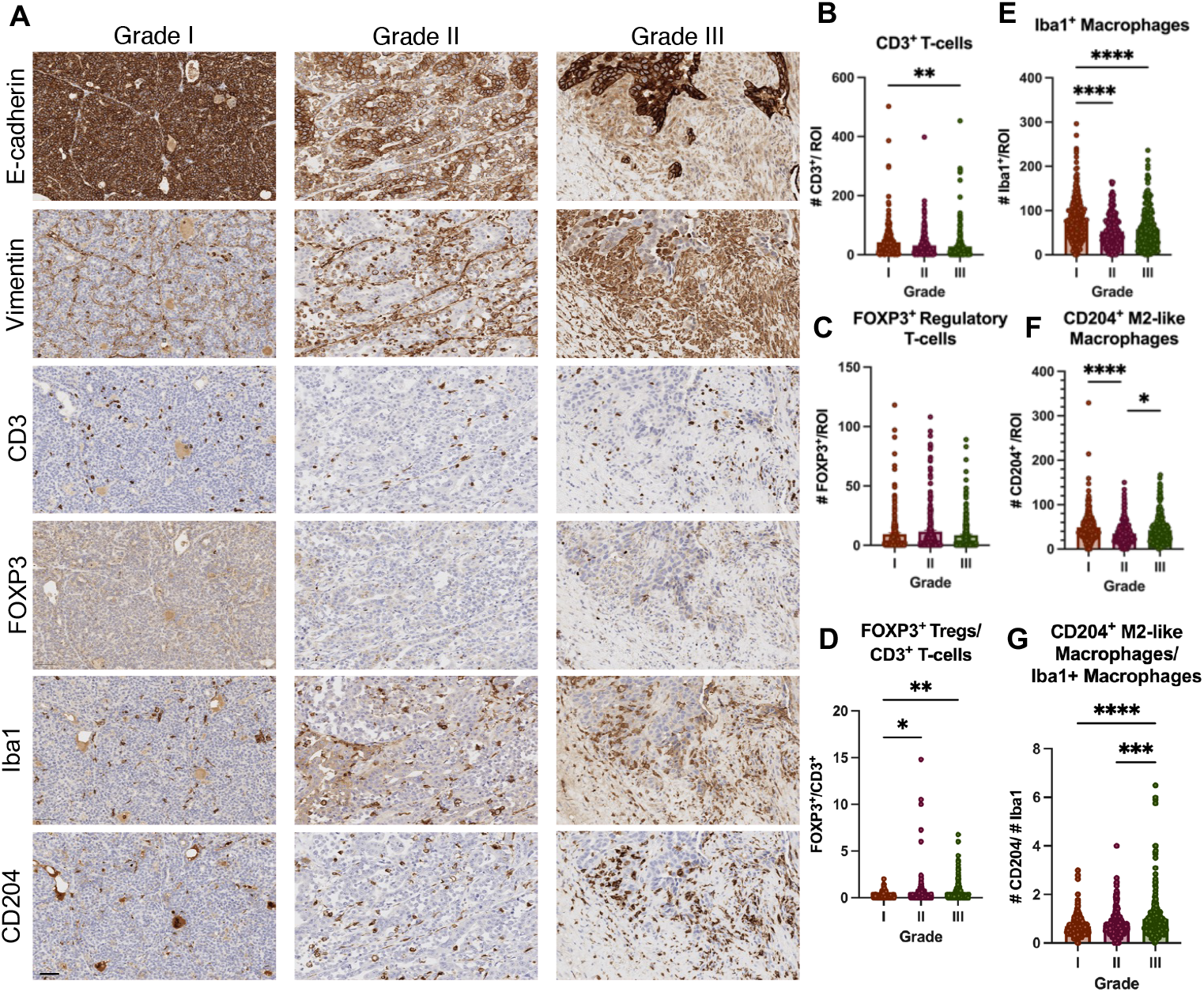
CMCs display an association between EMP and immune infiltration across histological grade. **(A)** Fifty-two archived FFPE CMC specimens were labeled for EMP (E-cadherin, Vimentin) and immune cell (CD3, FOXP3, Iba1, CD204) markers. **(B-G)** Immune infiltration was quantified across 15 high-power fields (HPF) for each tumor. All graphs are plotted as the mean value with error bars indicating the standard error of the mean. Data analyzed using one-way ANOVA: **** p < 0.0001, *** p < 0.001, ** p < 0.01, * p < 0.05, ns (p > 0.05) if not indicated. Scale bar indicates 50 µm. ROI = region of interest.

### Heterogeneous and quasi-mesenchymal CMCs assemble an immunosuppressive TME through upregulation of immunosuppressive paracrine factors

In the previous analysis, we identified a grade-based correlation between histologic grade, EMP status, and immune cell infiltration. To identify more specific associations between the EMP program and immunosuppression in CMCs, we selected a subset of 15 CMCs for additional histopathological analysis. In this subset of samples, we included 5 tumors of each of the following three categories: (i) epithelial (high E-cadherin expression and low vimentin expression in neoplastic cells, cuboidal to columnar cellular morphology), (ii) heterogeneous (tumors containing mixtures of epithelial and quasi-mesenchymal cells and cells residing in a partial state, co-expressing E-cadherin and vimentin and displaying a partial cellular morphology), and (iii) quasi-mesenchymal (low E-cadherin expression and high vimentin expression in neoplastic cells, partial and spindle-shaped cellular morphology). As expected, E-cadherin expression decreased and vimentin expression increased across this spectrum of EMP states, from epithelial to heterogeneous to quasi-mesenchymal **(Fig 3A-B)**. Additionally, the percentage of neoplastic cells co-expressing E-cadherin and vimentin was the highest in the heterogeneous tumors, on average displaying a ∼2.2x higher co-localization score compared to epithelial tumors and ∼1.4x higher compared to quasi-mesenchymal tumors. **(Fig 3C)**.

**Figure 3.**
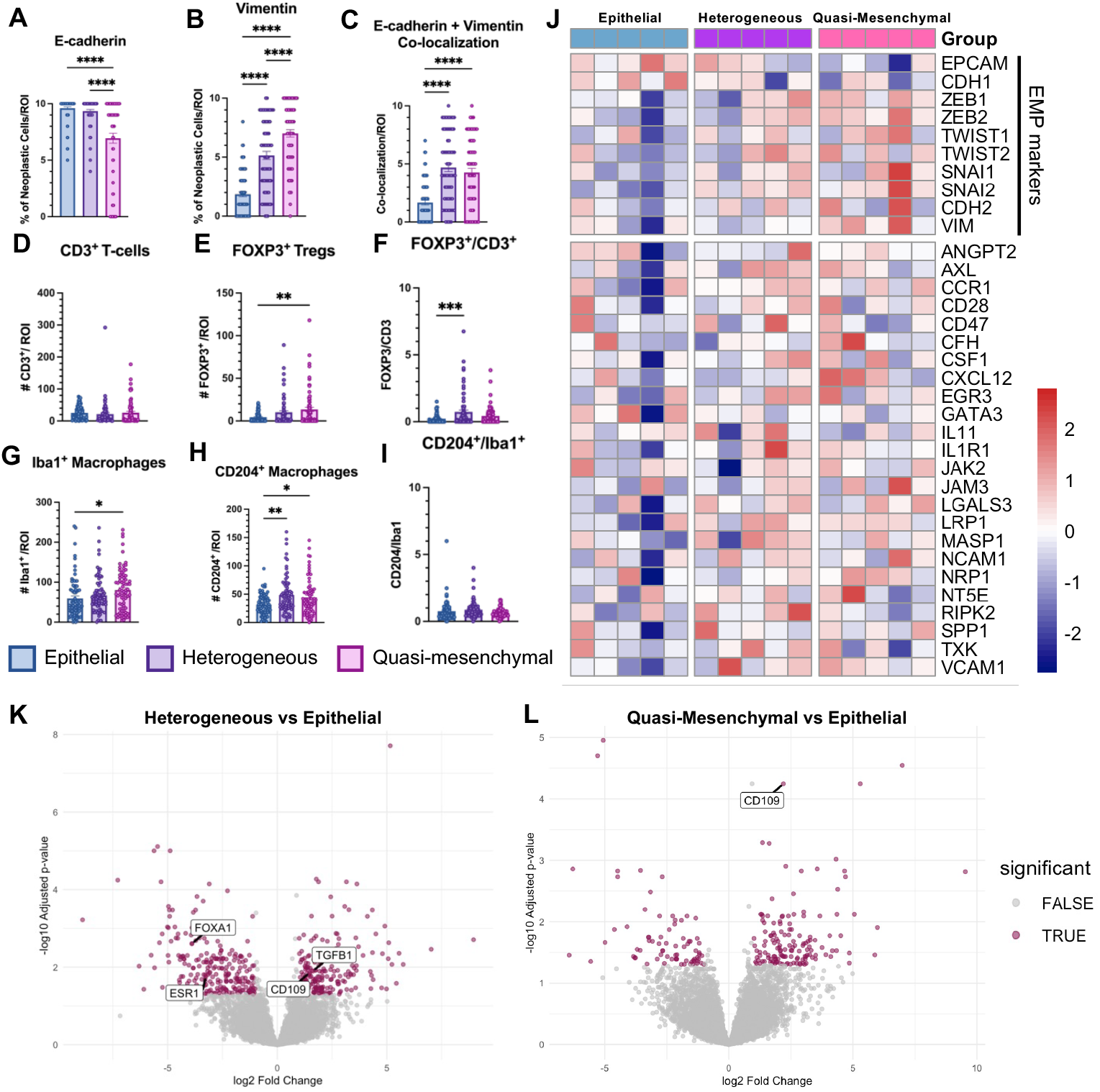
Heterogeneous and quasi-mesenchymal CMCs upregulate immunosuppressive paracrine factors. **(A-I)** Fifteen archived FFPE CMC specimens across epithelial, heterogeneous, and quasi-mesenchymal phenotypes were labeled for EMP (E-cadherin, vimentin) and immune (CD3, FOXP3, Iba1, CD204) markers by IHC and quantified across 15 high-power fields (HPF) for each tumor. All graphs are plotted as the mean value with error bars indicating the standard error of the mean. Data analyzed using one-way ANOVA: **** p < 0.0001, *** p < 0.001, ** p < 0.01, * p < 0.05, ns (p > 0.05) if not indicated. **(J)** The same 15 tumors were submitted for RNA sequencing (RNA-seq). The z-scores calculated from normalized expression of select EMP-related immunosuppressive factors are shown in the heatmap, along with the expression of several EMP markers. **(K-L)** Volcano plots displaying differentially-expressed genes between epithelial, heterogeneous, and quasi-mesenchymal tumors. Significantly differentially-expressed genes are defined as genes with abs(log_2_FC) > 1 and p_adj_ < 0.05. ROI = region of interest

Next, we measured immune infiltration across these three EMP states. Although we did not find differences in the number of CD3^+^ T-cells, we did observe that the number of infiltrating FOXP3^+^ Tregs increased in both the heterogeneous and quasi-mesenchymal tumors compared to epithelial tumors and the ratio of FOXP3^+^ Tregs/CD3^+^ T-cells was highest in the heterogeneous tumors **(Fig 3D-F)**. The number of Iba1^+^ macrophages and CD204^+^ M2-like macrophages was increased in both heterogeneous and quasi-mesenchymal tumors compared to epithelial tumors, however the ratio of CD204^+^ M2-like macrophages/Iba1^+^ macrophages was unchanged between these three states **(Fig 3G-I)**. Altogether, this suggests that both heterogeneous tumors containing combinations of epithelial, hybrid, and quasi-mesenchymal cells, as well as quasi-mesenchymal tumors recruit pro-tumor immune cells to the TME, including M2-like macrophages and Tregs. This is consistent with findings from our murine model, which have shown that mixed tumors containing both epithelial and quasi-mesenchymal cancer cells recruit Tregs and M2-like macrophages at similar numbers as compared to quasi-mesenchymal tumors (21).

To identify whether the heterogeneous and quasi-mesenchymal tumors upregulate immunosuppressive factors, leading to the formation of an immunosuppressive TME, we performed RNA-seq analysis on the same 15 tumors from Fig 3A-I. By principal component analysis (PCA), the epithelial samples showed greater separation from the heterogeneous and quasi-mesenchymal samples **(Fig S4A)**. Moreover, differential expression analysis of these three groups yielded many differentially expressed genes when comparing epithelial and either heterogeneous or quasi-mesenchymal samples, but not heterogeneous and quasi-mesenchymal samples **(Fig S4B-D)**. We first examined the expression of our previously-identified EMP-related paracrine factors (22). As expected, the heterogeneous and quasi-mesenchymal tumors downregulated the epithelial markers EPCAM and E-cadherin (*CDH1*) and instead upregulated several mesenchymal markers and EMT-TFs (ZEB1, ZEB2, TWIST1, TWIST2, Snail (*SNAI1*), Slug (*SNAI2*), N-cadherin (*CDH2*), and Vimentin (*VIM*) **(Fig 3J)**. The heterogeneous and quasi-mesenchymal tumors concomitantly upregulated several immunosuppressive genes. Of note, CSF1 and CXCL12 were upregulated in the quasi-mesenchymal tumors, whereas LGALS3, MASP1, CD73 (*NT5E*), and SPP1 were upregulated in both heterogeneous and quasi-mesenchymal tumors compared to epithelial tumors (**Fig 3J)**.

In addition to the upregSommer et al., 2021, Bladder Cancer (64). (D) Correlationulation of these immunosuppressive factors, we noted other differential expression patterns in our samples. The pro-inflammatory cytokine IFN-γ (*IFNG*) was downregulated in both the heterogeneous and quasi-mesenchymal tumors compared to epithelial tumors **(Fig S4G)**. Similarly, the heterogeneous and quasi-mesenchymal tumors also upregulated the immunosuppressive cytokine TGF-β1 (*TGFB1*) **(Fig 3K, S4I)**. Moreover, these heterogeneous and quasi-mesenchymal tumors showed increased expression of matrix metalloprotease MMP13 **(S4H)**. Upregulation of matrix metalloproteases are associated with the invasive features of EMP (8). Interestingly, the heterogeneous and quasi-mesenchymal tumors also downregulated estrogen receptor alpha (*ESR1*) compared to epithelial tumors **(Fig 3K, S4F)**. This is in concordance with findings that human triple-negative breast cancers (TNBC) activate the EMP program and downregulate hormone receptors, such as estrogen receptor (54, 55). Similarly, the epithelial CMCs showed increased expression of FOXA1, a transcription factor that is expressed by luminal breast cancer cell lines **(Fig 3K, S4E)**. Finally, we identified that the cell-surface glycoprotein CD109 was significantly upregulated in both heterogeneous and quasi-mesenchymal tumors compared to epithelial tumors **(Fig 3K-L)**. These findings revealed that EMP creates a TNBC-like phenotype in heterogeneous and quasi-mesenchymal tumors. Furthermore, we identified a potential mediator of EMP-driven immunosuppression: CD109.

### CD109 is a potential mediator of tumorigenicity and immunosuppression in quasi-mesenchymal breast cancers

The glycoprotein CD109 has been demonstrated to negatively regulate TGF-β signaling by acting as a co-receptor for this cytokine (56). CD109 has been associated with increased cell proliferation, migration, and stemness in several cancers including TNBC and squamous cell carcinoma (57– 59). Furthermore, CD109 overexpression reduced inflammation in a model of wound healing, but this immunosuppressive role has not been demonstrated in the neoplastic setting (60). These previous studies established that CD109 has independent roles in cancer progression and immunosuppression, but these had previously not been associated together, and particularly not within the context of EMP. Therefore, we aimed to elucidate the associations between CD109 and EMP-mediated immunosuppression in both CMCs, as well as human breast cancers.

CD109 was significantly upregulated in quasi-mesenchymal CMCs compared to epithelial CMCs (4.6x higher mean expression), with intermediate expression in heterogeneous tumors (**Fig 4A)**. Using a published RNA-seq dataset of 197 naturally-occurring CMCs (61), we found that the expression of CD109 negatively correlated with epithelial markers EPCAM and E-cadherin (*CDH1*) and positively correlated with EMT-TFs ZEB1, ZEB2, TWIST1, Snail (*SNAI1*), Slug (*SNAI2*), as well as the mesenchymal marker vimentin (*VIM)* (**Fig 4B-C)** (61). Using histologic and transcriptomic sample information, we found that CD109 expression positively correlated with histologic grade and was highly expressed in basal-like CMCs **(Fig 4D-E)**. Finally, dogs with CMCs expressing high levels of CD109 showed a trend towards decreased survival (p = 0.1844) compared to dogs with CMCs expressing low levels of CD109 **(Fig 4F)**.

**Figure 4.**
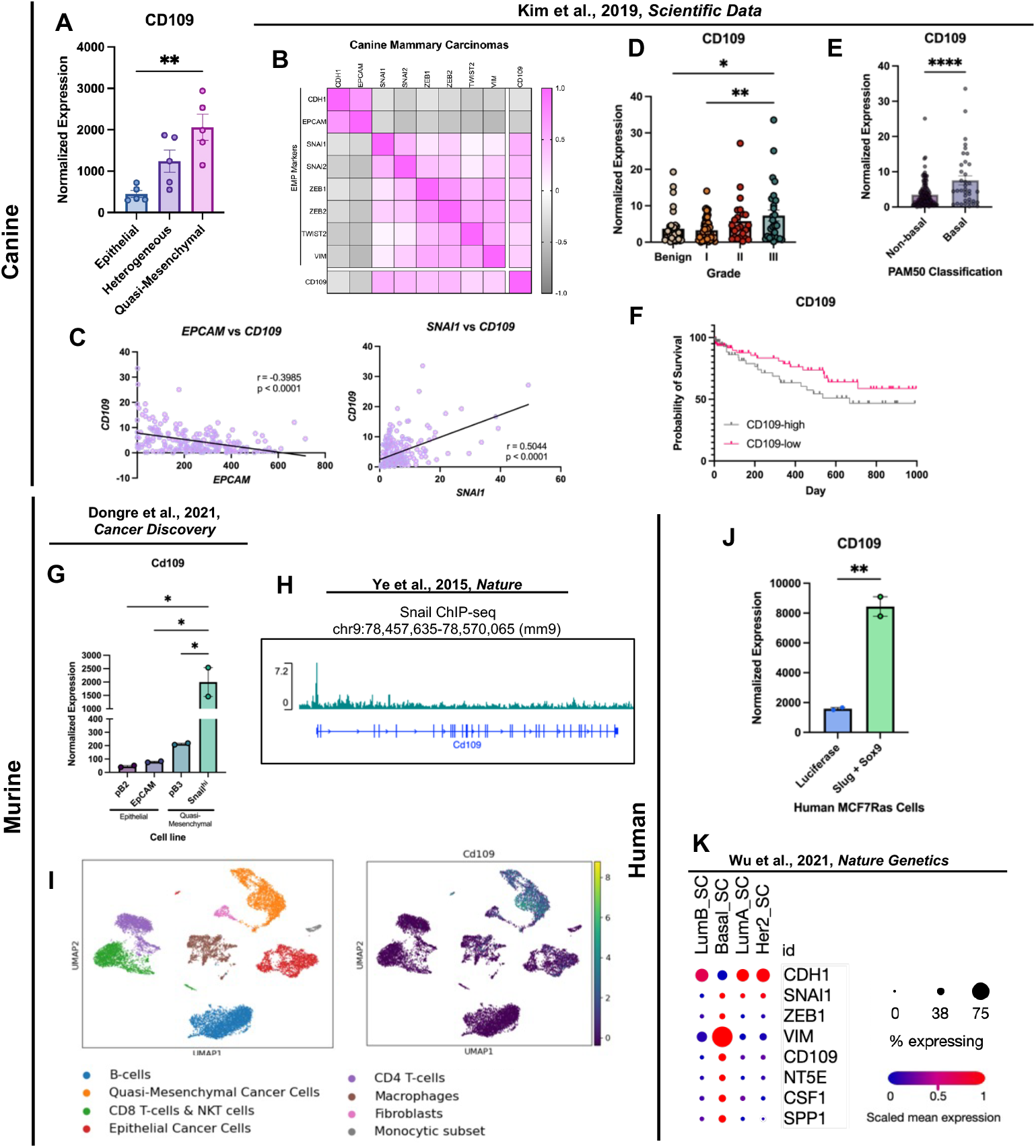
EMP is associated with increased CD109 expression in canine, murine, and human breast carcinomas. **(A)** Normalized expression of CD109 in CMCs of increasing EMP from RNA-seq data presented in Fig 3. **(B)** Correlation matrix demonstrating association between CD109 expression and EMP markers in CMCs. Scale bar indicates Pearson correlation coefficients. **(C)** Pearson correlation analysis demonstrating anticorrelation of EPCAM and CD109 and correlation of Snail and CD109. **(D)** Normalized CD109 expression across CMC histopathological grade. **(E)** CD109 expression in non-basal and basal CMCs, based on PAM50 classification. **(F)** Survival curve of dogs bearing CMCs with high or low CD109 expression. Data analyzed using Mantel-Cox Log-Rank test (p = 0.1844). **(B-F)** Data from Kim et al., 2019, *Scientific Data* (61). **(G)** Normalized expression of CD109 in epithelial and quasi-mesenchymal murine mammary tumor cell lines. Data from Dongre et al., 2021, *Cancer Discovery*. **(H)** Chromatin immunoprecipitation sequencing (ChIP-seq) data demonstrating peaks for Snail binding around the Cd109 gene. Data from Ye et al., 2015, *Nature* (6). **(I)** Single-cell RNA sequencing (scRNA-seq) data of heterogeneous murine mammary tumors demonstrates that CD109 expression is mainly localized to quasi-mesenchymal cancer cells. **(J)** Upon doxycycline-controlled overexpression of EMT-TFs Slug and Sox9, human MCF7RAS breast cancer cells upregulate CD109 compared to control luciferase-expressing cells. Data from Dongre et al., 2021, *Cancer Discovery* (22). **(K)** scRNA-seq of human breast primary tumors reveals that basal-like epithelial cancer cells upregulate CD109, along with EMT-TFs and immunosuppressive factors. Data from Wu et al., 2021, *Nature Genetics* (62). All bar graphs are plotted as the mean value with error bars indicating the standard error of the mean. Data analyzed using one-way ANOVA (A, C, F) or student’s t-test (D, I): **** p < 0.0001, *** p < 0.001, ** p < 0.01, * p < 0.05, ns (p > 0.05) if not indicated.

We next sought to determine whether the association between EMP and CD109 expression could be translated to models in other species. Using our MMTV-PyMT based murine model of epithelial and quasi-mesenchymal mammary carcinomas, we found that the epithelial mammary tumor cell lines (pB2, EpCAM) express low levels of CD109 compared to the quasi-mesenchymal mammary tumor cell lines (pB3, Snail^hi^) **(Fig 4G)**. The Snail^hi^ cell line contains cells sorted based on high expression of the EMT-TF Snail and showed significantly higher CD109 expression, suggesting an even greater association between EMP and CD109 expression. To identify whether CD109 is directly regulated by the EMP program, we analyzed publicly-available Chromatin immunoprecipitation sequencing (ChIP-seq) data for the EMT-TF Snail in pB3 quasi-mesenchymal murine mammary cancer cells (6). We found that the *Cd109* gene contains several binding sites for Snail, particularly near the *Cd109* transcription start site, demonstrating that it is directly regulated by the EMT-TF Snail **(Fig 4H)**.

To confirm that quasi-mesenchymal cancer cells are the predominant source of CD109 in murine mammary tumors, we performed single-cell RNA sequencing (scRNA-seq) of a mixed epithelial-quasi-mesenchymal tumor. To generate this artificially heterogeneous tumor, we implanted a 9:1 mixture of epithelial and quasi-mesenchymal MMTV-PyMT-derived murine mammary cancer cells into the mammary fat pad of syngeneic hosts. Using this model, we identified that CD109 expression is localized mainly to quasi-mesenchymal cancer cells within heterogeneous murine MMTV-PyMT-derived tumors, with few epithelial cells and macrophages also showing low CD109 expression **(Fig 4I)**.

Finally, we wanted to identify whether the association between EMP and CD109 expression translates to human breast cancers. We first used MCF7Ras human breast cancer cells with doxycycline-controlled expression of the EMT-TFs SNAI2 and SOX9 (3, 22, 55). Upon doxycycline treatment, these cells activate EMP and adopt a quasi-mesenchymal morphology, lose expression of E-cadherin, and upregulate vimentin (55). Analysis of RNA-seq data obtained from SNAI2 and SOX9-induced MCF7Ras cells compared to control cells with doxycycline-controlled expression of luciferase show a four-fold increase in CD109 expression **(Fig 4J)**. Additionally, analysis of scRNA-seq data from human primary breast tumors showed that epithelial cells with basal-like expression upregulate CD109, as well as other immunosuppressive factors including CD73 (*NT5E*), CSF1, and SPP1 **(Fig 4K)** (62). Altogether, these findings demonstrate that CD109 is directly upregulated by EMP and correlates with immune-evasive features of quasi-mesenchymal cancer cells in multiple species.

### Integration of transcriptomic data across canine carcinoma datasets reveals conserved drivers of EMP-mediated immunosuppression

After determining that EMP is associated with immunosuppression in CMCs, we sought to identify if this process was conserved across other canine cancers of epithelial origin. We explored published RNA sequencing (RNA-seq) data from CMCs (n=197) (61), canine oral squamous cell carcinoma (n=8) (63), canine invasive urothelial carcinoma (n=56) (64), and canine pulmonary carcinoma (n=12) (65) to measure associations between the expression of EMP markers and immunosuppressive genes. For each dataset, we first filtered the genes to only include EMP-related paracrine factors. Several of these EMP-related paracrine factors have been described to be important for tumor progression and immune evasion in murine models but have yet to be associated with EMP and immunosuppression in naturally-occurring canine tumors. To address this, we calculated Pearson correlation coefficients for the normalized expression of selected EMP markers and each EMP-related paracrine factor in all four tumor types.

In CMC samples, we found 13 EMP-related paracrine factors to be positively correlated with mesenchymal markers (vimentin, *VIM*) and EMT-TFs (Snail (*SNAI1*), Slug (*SNAI2*), ZEB1, ZEB2, TWIST2) and negatively correlated with epithelial markers (E-cadherin (*CDH1*) and EPCAM). These factors included LRP1, IL1R1, CD73 (*NT5E*), NRP1, AXL, CXCL12, ANGPT2, JAM3, CFH, CCR1, MASP1, CSF1, and VCAM1 **(Fig 5A)**. We were particularly intrigued to see CD73, MASP1, CXCL12, and CSF1 show correlations with EMP as these factors were also identified to have immunosuppressive functions in our previously published murine model (21, 22). Specifically, knockout of CD73 and CSF1 from quasi-mesenchymal murine mammary carcinoma cells remodeled the TME to allow infiltration of anti-tumor T-cells and M1-like macrophages and at least partially sensitized quasi-mesenchymal tumors to anti-CTLA-4 ICB therapy (22). In oral squamous cell carcinoma samples, SPP1, CXCL12, VCAM1, NRP1, and CD73 were correlated with mesenchymal markers (N-cadherin (*NCAD*), vimentin (*VIM*)) and EMT-TFs (Snail (*SNAI1*), ZEB1, and TWIST2) but not epithelial markers (E-cadherin (*CDH1*) and EPCAM) **(Fig 5B)**. Again, SPP1, CXCL12, and CD73 were previously identified to be associated with immunosuppressive functions in our murine mammary tumor model. Furthermore, targeting SPP1 alters the TME and partially sensitizes quasi-mesenchymal tumors to anti-CTLA-4 (22). In invasive urothelial carcinoma samples, NCAM1, CXCL12, and SPP1 were positively correlated with EMT-TF ZEB1 and mesenchymal marker vimentin (*VIM*) and negatively correlated with epithelial markers E-cadherin (*CDH1*) and EPCAM **(Fig 5C)**. Finally, pulmonary carcinoma samples showed positive correlations between CFH, CD73 (*NT5E*), SPP1, AXL, and ANGPT2 with mesenchymal marker Vimentin (*VIM*) and EMT-TFs Snail (*SNAI1*), ZEB1, ZEB2, and TWIST2, but not epithelial markers E-cadherin (*CDH1*) and EPCAM **(Fig 5D)**.

**Figure 5.**
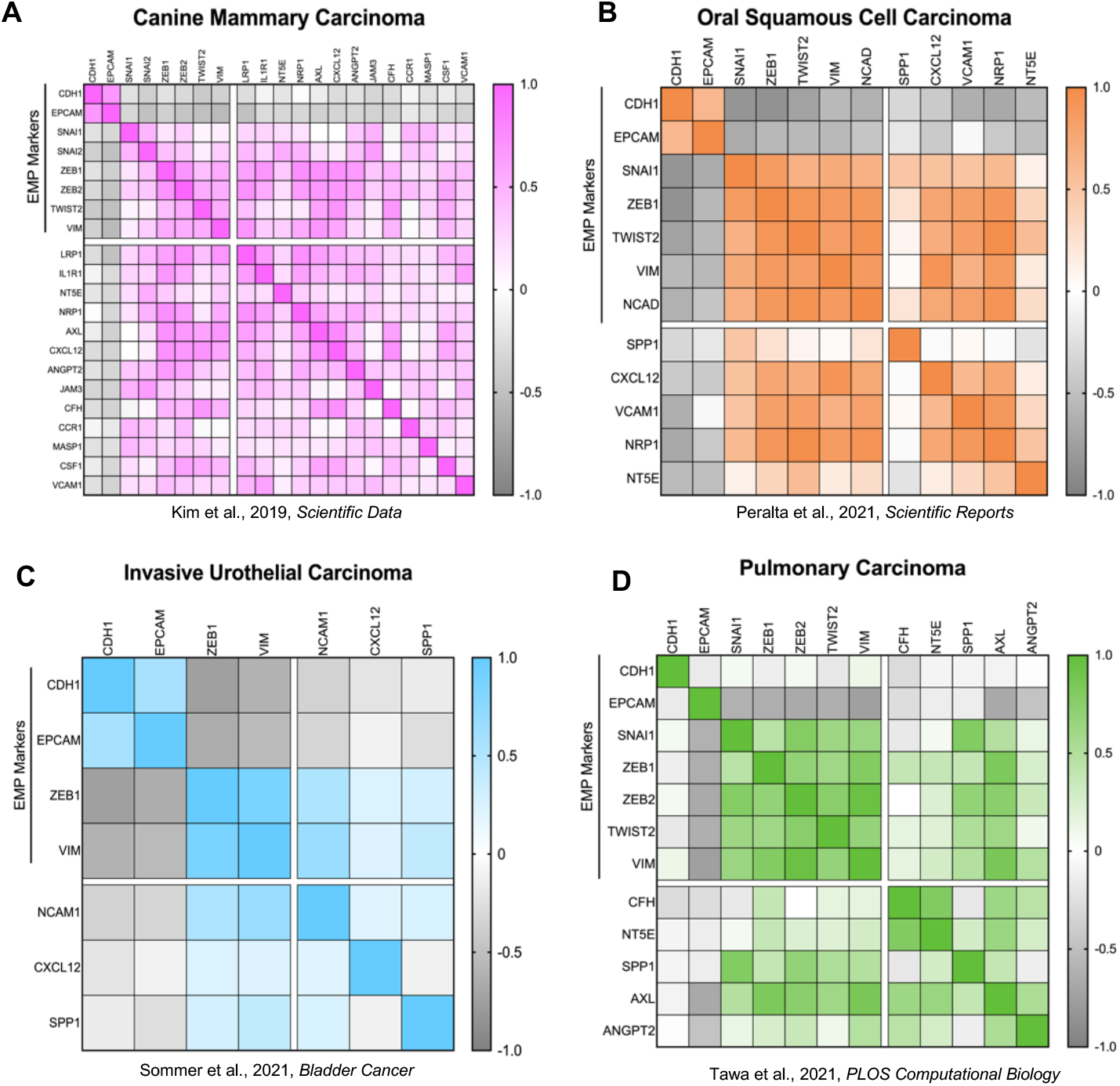
EMP markers correlate with expression of immunosuppressive paracrine factors across multiple canine cancer types. **(A)** Correlation matrix showing EMP-associated immunosuppressive genes (Dongre et al., 2021, *Cancer Discovery;* 22) that correlate with expression of EMT-TFs and mesenchymal markers in CMCs. Data from Kim et al., 2019, *Scientific Data* (61). **(B)** Genes that were positively correlated with expression of EMT-TFs and mesenchymal markers in oral squamous cell carcinoma samples are indicated. Data from Peralta et al., 2021, *Scientific Reports* (63). **(C)** Genes that were positively correlated with expression of EMT-TFs and mesenchymal markers in invasive urothelial carcinoma samples are shown in the correlation matrix. Data from Sommer et al., 2021, *Bladder Cancer* (64). **(D)** Correlation matrix showing genes that positively correlated with expression of EMT-TFs and mesenchymal markers in pulmonary carcinoma samples. Data from Tawa et al., 2021, *PLOS Computational Biology* (65). Scale bars indicate Pearson correlation coefficients.

This analysis of publicly available transcriptomics data from several canine cancers allowed us to identify that EMP is associated with upregulation of immunosuppressive factors in four distinct carcinomas. We focused on the EMP-related paracrine factors because these have been demonstrated to have immunosuppressive roles in murine models. Among these four cancers, CD73, SPP1, and CXCL12 were associated with EMP markers in three cancers. Furthermore, ANGPT2, CFH, AXL, NRP1, and VCAM1 were correlated with EMP markers in two tumor types. The upregulation of these immunosuppressive factors in tumor samples with higher expression of mesenchymal markers and EMT-TFs suggests that these may be conserved drivers of EMP-mediated immunosuppression and resistance to anti-tumor immunity in multiple cancers.

## Discussion

EMP is well-established as a driver of invasion, metastasis, and chemotherapy resistance. Recent studies by us and others demonstrated that EMP also drives immunosuppression in murine cancer models, particularly breast cancer, hepatocellular carcinoma, pulmonary carcinoma, and melanoma (21, 23–25, 50, 53). However, this trend had previously not been demonstrated in naturally-occurring canine carcinomas. We chose to investigate EMP-mediated immunosuppression in spontaneous CMCs due to their similarities to human breast cancers in terms of heterogeneity, clinical features, and biological pathways. We found that not only CMCs, but several other naturally occurring canine carcinomas including oral squamous cell carcinoma, invasive urothelial carcinoma, and pulmonary carcinoma upregulate immunosuppressive genes, which correlate with EMP markers. In CMCs, the expression of these immunosuppressive factors correlated with a shift of the TME from immunopermissive to immunosuppressive. By comparing CMCs spanning several EMP states, we identified a connection between EMP and CD109 expression. This glycoprotein is somewhat poorly characterized and its relationship to EMP is unclear. Furthermore, CD109 expression had never been associated with an immunosuppressive TME. We were able to demonstrate that EMP is associated with CD109 expression in three species, highlighting its translational relevance as a therapeutic target and biomarker. Our study sheds light on the ability of the EMP program to drive immunosuppression in multiple naturally-occurring cancers and we propose that identifying and functionally testing EMP-regulated immunosuppressive genes like CD109 may be useful in sensitizing heterogeneous and quasi-mesenchymal tumors to immune attack in multiple species.

We first demonstrate that high-grade CMCs activate EMP and this is associated with the assembly of an immunosuppressive TME. The grade III CMCs displayed higher expression of the mesenchymal marker vimentin along with cytoplasmic translocation of E-cadherin and a de-differentiated, partial cellular morphology. Moreover, these high-grade tumors showed increased percentages of neoplastic cells co-expressing E-cadherin and vimentin. This is consistent with previous observations that upon EMP activation *in vivo*, cells tend to reside in a partial or quasi-mesenchymal state, rather than transitioning to a completely mesenchymal phenotype. Furthermore, cells residing in these partial states tend to be more invasive than those in the extreme mesenchymal state (18–20). Of note, the apparent percentage of cells co-expressing E-cadherin and vimentin by IHC may have been inflated by the presence of well-differentiated myoepithelial cells within these lower-grade tumors. Neoplastic myoepithelial cells replicate the physiological patterns of membranous E-cadherin labeling and cytoplasmic vimentin labeling observed in healthy mammary glands. Grade I/III and II/III carcinomas tend to retain a higher degree of ductal and glandular differentiation and therefore contain comparatively increased numbers of these cells. Nonetheless, we observed that the grade III CMCs also recruited greater proportions of pro-tumor immune cells. Notably, heterogeneous tumors containing mixtures of epithelial, hybrid, and quasi-mesenchymal cancer cells recruited high numbers of M2-like macrophages and an increased proportion of Tregs/T-cells, suggesting that a fraction of hybrid or quasi-mesenchymal cancer cells may be sufficient to drive resistance to immune attack. These findings closely match our previous murine studies that demonstrate that quasi-mesenchymal cancer cells, even in minority fractions, drive resistance to anti-tumor immunity.

In addition to the assembly of an immunosuppressive TME, these heterogeneous and quasi-mesenchymal CMCs also upregulated several EMP-related paracrine factors. This is intriguing for two reasons. Firstly, these factors were found to be associated with EMP-mediated immunosuppression in a completely different species. Secondly, several of these factors were upregulated in both heterogeneous and quasi-mesenchymal CMCs. Once again, this suggests that a fraction of hybrid or quasi-mesenchymal cancer cells can lead to an increase in immunosuppressive factors within CMCs. However, a limitation of this study is the use of bulk RNA-seq, which does not allow us to pinpoint the source of these immunosuppressive factors. Future studies should use approaches like scRNA-seq and spatial transcriptomics to determine which cells are producing these factors and where in the tumor they are located.

We were particularly interested to observe that these cancers upregulated CD73 (*NT5E)*, CSF1, and SPP1, which have been associated with immunosuppressive functions in our previous work (22). Specifically, our group and others have identified that targeting CD73 sensitizes murine breast cancer and colon cancer models to anti-CTLA-4 and anti-PD-1 immune checkpoint blockade therapies (22, 55, 66). Moreover, EMP is associated with CD73 expression in human breast cancer (55, 67). CSF1/CSF1R blockade enhances anti-CTLA-4 and anti-PD-1 efficacy in a murine orthotopic pancreatic adenocarcinoma model (68). And finally, targeting SPP1 reduces tumor growth in a murine colorectal cancer model (69). Our findings in naturally-occurring CMCs therefore provide rationale for targeting these immunosuppressive factors in heterogeneous and quasi-mesenchymal human tumors. Furthermore, it suggests that the mechanisms of EMP-mediated immunosuppression in carcinomas translates across species from orthotopic murine tumors to naturally-occurring canine tumors and potentially even human cancers.

In addition to these established immunosuppressive factors, we identified that CD109 associates with EMP and may play an immunosuppressive role in heterogeneous and quasi-mesenchymal tumors. CD109 has been associated with tumorigenicity of squamous cell carcinoma and TNBC, but has a contested association with EMP. Here, we show that EMP is correlated with increased CD109 expression across three species: murine, canine, and human. Additionally, CD109 has been shown to dampen inflammation in wound healing, but this immunosuppression has not been associated with cancer. Our findings indicate that EMP, tumorigenicity, and immunosuppression may all be associated with CD109 activity. Interestingly, EMP is activated during both wound healing and cancer progression, suggesting that EMP-induced CD109 expression may be observed in several biological contexts. Although we demonstrated an association, further work is needed to elucidate the functional role for CD109-mediated immunosuppression. Moreover, the high expression of both EMP-inducing factor TGF-β and its negative regulator CD109 in quasi-mesenchymal cancer cells is somewhat paradoxical and warrants further investigation. This co-expression suggests that factors other than TGF-β may be sufficient to induce EMP in these tumors. Finally, it is unknown whether the non-cancer cells express CD109 in CMCs and where in these heterogeneous tumors CD109 is expressed. Therefore, mapping the spatial location of CD109 expression in comparison to regions of quasi-mesenchymal cancer cells will also provide valuable insight to the tumorigenic functions of CD109.

Finally, we show that the association of EMP and immunosuppression is observed in three additional canine tumor types. Certain factors like CD73, SPP1, and CXCL12 correlated with expression of EMP markers across the four canine cancers (CMC, oral squamous cell carcinoma, invasive urothelial carcinoma, and pulmonary carcinoma). Other factors appeared to display cancer-specific associations with EMP. Interestingly, the correlations between CD109 expression and EMP was observed in CMCs, but not the other three canine cancers. These findings suggest that EMP-mediated immunosuppression is observed in several different naturally-occurring canine carcinomas, although the identity of the immunosuppressive factors may differ. The reason for these differences is unknown but may suggest that the cell of origin and its microenvironment regulate the expression of immunosuppressive factors.

In sum, this study highlights the use of naturally-occurring canine tumors to substantiate findings from murine models and provide translational relevance for human cancers. Here, we used this translational framework to identify pan-cancer (CD73, SPP1, and CXCL12) as well as CMC-specific (CD109) associations between EMP and immunosuppression. These factors hold the potential to be useful as biomarkers and therapeutic targets to sensitize immune-resistant tumors to immunotherapies in both canine and human cancer patients.

## Materials and Methods

### Sample selection and histologic analysis

A total of 52 formalin-fixed, paraffin-embedded (FFPE) CMC sections from 51 canine mammary carcinoma cases were retrieved from the archives of the New York State Veterinary Diagnostic Laboratory, Cornell University College of Veterinary Medicine, Section of Anatomic Pathology (Table S1). Cases were identified in our internal institutional software by filtering on the species (“Canine”) and using the terms “mammary carcinoma”, AND/OR “carcinoma”, AND/OR “mammary”, AND/OR “grade I/grade 1” AND/OR “grade II/grade 2” AND/OR “grade III/grade 3”. Seventeen cases were retrieved per grade (17 grade I out of III, 17 grade II out of III, 17 grade III out of III). Upon regrading of the CMC samples, there were 14 grade I out of III, 18 grade II out of III, and 20 grade III out of III tumors. Because surgically excised CMCs are often either a large mass or excised together with part of or the entire mammary chain, adequate histologic evaluation typically requires between 1-15 slides to be prepared on average. A single representative section was thus selected per case, ensuring that: (**i**) the histomorphologic features of cancer cells were representative of other sections taken from the same mass; (**ii**) the section included the pre-existing cutaneous tissue when available (deep and lateral surgical margins); (**iii**) the degree of immune infiltration (periphery and center) was representative of other sections taken from the same mass; (**iv**) any other histomorphologic features (e.g. vascular invasion, necrosis) were adequately represented compared to other sections.

CMC samples were reviewed by two board-certified veterinary anatomic pathologists (SN & ADM). CMCs were blindly re-evaluated for subtype and grade (SN). Where discrepancies with the original diagnosis arose (SN evaluation), cases were further reviewed by ADM to ensure a tri-partite review of complex cases. CMCs were graded using the criteria defined by Peña et al. that are routinely used in diagnostic pathology (70): (i) percent tubule formation: > 75%, 10-75%, or < 10%; (ii) nuclear pleomorphism: mild pleomorphism and uniform/small nuclei with occasional nucleoli; moderate anisokaryosis and nuclear morphological variation with increased nucleoli; marked nuclear pleomorphism with hyperchromasia and prominent nucleoli and (iii): mitotic activity: 0-9 mitoses/10 HPF; 10-19 mitoses/10 HPF; > 20 mitoses/10 HPF (48, 70).

Cancer cells were first categorized based on their morphology using routine hematoxylin-eosin staining: (**i**) overtly epithelial (cuboidal, columnar, squamous), (**ii**) overtly mesenchymal (elongated and spindle-shaped with a central nucleus and either acicular ends or double-blunted ends), and (**iii**) partial/hybrid (morphology that does not fall under either epithelial or mesenchymal category, *e*.*g*. plump cells with a pinched acicular base and plump rounded apex). Histomorphologic characteristics were then compared to labeling patterns of E-cadherin and vimentin: overtly cytoplasmic, overtly membranous, and myoepithelial-like (vimentin: moderate to marked cytoplasmic labeling intensity; E-cadherin: moderate to marked membranous cytoplasmic labeling intensity).

### Immunohistochemical labeling and quantification

Immunohistochemistry (IHC) was used to label each sample for the following markers: E-cadherin (clone DECMA-1, Millipore #MABT26, RRID: AB_10807576), vimentin (clone Vim 3B4, Dako #M7020), CD3 (clone LN10, Leica #PA0553, RRID: AB_10554601), FOXP3 (clone FJK-16s, Invitrogen #14-5773-82), Iba1 (polyclonal, Wako #019-19741, RRID: AB_839504), CD204 (clone SRA-E5, Cosmo Bio # KAL-KT022). IHC staining was conducted as previously described (71, 72).

IHC and H&E slides were scanned at 40x using the Aperio CS2 ScanScope Unit provided by the Cornell University Department of Biomedical Sciences. IHC scoring and immune cell quantification was performed using QuPath (73). For E-cadherin and vimentin labeling, 15 randomly selected regions of interest (ROIs) were scored for % of cells expressing the marker (0-10 scale), staining intensity (0-3 scale), and notes about localization of the marker as well as morphology of the stained cells were recorded. The two numeric scores (% of cells expressing the marker and staining intensity) were then multiplied to generate a composite score. Co-localization of E-cadherin and vimentin labeling was estimated by visual assessment of matched ROIs for each tumor. For CD3, FOXP3, Iba1, and CD204, the number of positively labeled cells were quantified across 15 randomly selected ROIs. The standard dimension for each ROI was 490µm x 490µm, which represents one digital high-power field (10 high-power fields: 2.37mm^2^). The ROIs were quantified/scored independently by two veterinary students (KB, LL) and a veterinary anatomic pathologist (SN).

### CMC RNA extraction, sequencing, and analysis

Adjacent normal tissue was trimmed from blocks prior to sectioning for RNA-seq. One 10 mm scroll was then cut from each of fifteen CMC formalin-fixed paraffin-embedded (FFPE) samples and submitted for RNA extraction and sequencing. All RNA extraction, library preparation, sequencing, and gene expression analysis was performed by the Cornell University Biotechnology Resource Center Transcriptional Regulation and Expression Facility (RRID: SCR_022532).

RNA was extracted using the FormaPure XL RNA Reagent Kit (Beckman Coulter #C36000). The FormaPure XL RNA Reagent Kit was used following the manufacturer’s instructions, except that samples were incubated with proteinase K overnight, with gentle shaking. RNA sample quality was confirmed by spectrophotometry (Nanodrop) to determine concentration and chemical purity (A260/230 and A260/280 ratios) and with a Fragment Analyzer (Agilent) to determine RNA integrity. For RNAseq, 750ng total RNA per sample was treated with the FastSelect rRNA depletion kit (Qiagen) to remove rRNA. UDI-barcoded RNAseq libraries were generated with the NEBNext Ultra II Directional RNA Library Prep Kit (New England Biolabs). Each library was quantified with a Qubit (dsDNA HS kit; Thermo Fisher) and the size distribution was determined with a Fragment Analyzer (Agilent) prior to pooling. Libraries were sequenced on a NovaSeqX instrument (Illumina) to generate 2×150 PE reads. At least 20M read pairs were generated per library.

Reads were trimmed for low quality and adaptor sequences with fastp v0.21.0 (74). Reads were mapped to the reference genome/transcriptome (Ensembl CanFam4) using STAR v2.7.0e (75). SARTools and DESeq2 v1.26.0 was used to generate normalized counts and statistical analysis of differential gene expression (76, 77).

### Transcriptomic and chromatin immunoprecipitation sequencing (ChIP-seq) analysis of murine and human epithelial and quasi-mesenchymal cell lines

RNA-seq data of murine epithelial and quasi-mesenchymal MMTV-PyMT tumor-derived cell lines as well as MCF7Ras luciferase and Slug/Sox9-overexpressing human breast cancer cells have been previously published and are available through the Gene Expression Omnibus (GEO) database under accession code GSE161748 (22). ChIP-seq data are also previously published and available through GEO using accession number GSE61198 (6). ChIP-seq data (SRR1569438) were aligned to the mm9 mouse reference genome using bowtie2 with default settings. SAM files were converted to BAM format, sorted, and indexed using SAMtools. To generate signal intensity tracks, genome-wide read coverage was calculated from the sorted BAM files using deepTools. The resulting BigWig file was visualized using the Integrative Genomics Viewer (78).

### Mice

C57BL/6J female mice, aged 6–8-weeks, were obtained from The Jackson Laboratory. All animal procedures were carried out in compliance with guidelines and protocols approved by the Institutional Animal Care and Use Committee (IACUC) and maintained by the Center for Animal Resources and Education (CARE) at Cornell University.

### Generation and processing of murine mixed tumors for single-cell RNA sequencing (scRNA-seq)

To generate mixed epithelial-quasi-mesenchymal murine tumors, 1 x 10^6^ cells containing a 9:1 mixture of murine epithelial (pB2) and quasi-mesenchymal (Snail^hi^) cells were resuspended in 30 µl of media containing 20% Matrigel. Cells were implanted orthotopically into the mammary fat pad of C57BL/6J mice. After the tumors reached 2000 mm^3^, mice were euthanized, and tumors were excised. To generate single-cell suspensions, the tumors were first minced using a razor blade and digested in RPMI containing 2 mg/ml Collagenase A (Krackeler Scientific) and 100 U/ml Hyaluronidase (Krackeler Scientific). The suspension was incubated in a rotator at 37°C for 40 mins. Following digestion, the single-cell suspension was filtered through a 70-micron filter, followed by a 40-micron pore-sized strainer. The samples were then centrifuged at 1250 rpm for 10 mins at 4°C. The samples were resuspended in 1 mL of ACK Lysing Buffer (Gibco) and incubated at room temperature for 5 minutes to allow for hypotonic lysis of red blood cells. After 5 minutes, 1 mL warm RPMI was added, and the samples were centrifuged at 1250 rpm for 10 minutes. The pellet was then resuspended in Dead Cell Removal MicroBeads (Miltenyi Biotec) and incubated at room temperature for 15 minutes. After incubation, 500 µL of 1X binding buffer (Miltenyi Biotec) was added and the suspension was placed on a magnet stand for 5 minutes to remove dead cells. The supernatant was then transferred to a new tube and centrifuged at 1250 rpm for 5 minutes. The pellet was resuspended in 6200 µL cold PBS (Gibco) and 1800 µL Debris Removal Solution (Miltenyi Biotec) was mixed with the PBS. Then, 4 mL of cold PBS was overlayed on this mixture, and the tubes were centrifuged at 3000xg for 10 minutes at 4°C. The top two phases were discarded from the tube and 15 mL cold PBS was added to the bottom phase. The tubes were inverted three times and then centrifuged for 10 minutes at 1000xg at 4°C. Finally, the pellet was resuspended in PBS containing 0.04% Bovine Serum Albumin (BSA, Sigma-Aldrich) at a final concentration of 1,000 cells/µl. Single cell suspensions were run on a Chromium X instrument and libraries were prepared following the Chromium GEM-X Single Cell 3’ v4 Assay (10x Genomics, user guide CG000731, RevA) by the Cornell BRC Genomics Facility (RRID:SCR_021727). We targeted 10,000 cells per sample and used 12 cycles of cDNA amplification. Sample quality was confirmed using a Qubit (DNA HS kit; Thermo Fisher) to determine concentration and a Fragment Analyzer (Agilent) to confirm fragment size integrity. Libraries were sequenced on a NovaSeqX, 10B flowcell with 2 x 150bp read length.

### scRNA-seq analysis of murine mixed tumors and human breast primary tumors

scRNA-seq data from murine mixed tumors was aligned, filtered, and counted using Cell Ranger. Scanpy was used to exclude cells with less than 1000 counts or 200 genes, as well as cells with more than 30% of counts corresponding to mitochondrial genes. Genes expressed in less than 4 cells were also excluded. Counts were normalized and cells were clustered using the Louvain method. Cluster identities were assigned based on expression of the following markers: epithelial (Epcam, Cdh1, Tjp1, Itga6), quasi-mesenchymal (Snai1, Vim, Zeb1, Cdh2), CD4 T-cells (Ptprc, Cd3d, Cd3e, Cd3g, Tcf7, Cd28, Trbc2, Cd4), CD8 T-cells (Ptprc, Cd3dd, Cd3e, Cd3g, Tcf7, Cd28, Trbc2, Cd8a), B-cells (Ptprc, Cd19, Ms4a1, Cd79a, Pou2af1, Cd86), Macrophages (Ptprc, Itgam, Adre1, Csf1r, Cd68, Cd86, Fcgr3, Cd74, Mrc1, Arg1), Fibroblasts (Dcn, Col3a1, Col1a2, Col1a1, Pdgfra), Monocytic subset (Ptprc, Itgam, Cd14, Cd84, Arg2, Ly6gc1, Nos2).

Publicly-available scRNA-seq data of 26 human breast primary tumors were accessed through the Single Cell Portal (79). Samples were filtered to only include epithelial cells and EMP markers and immunosuppressive factors were selected to correlate expression.

### Cell lines and cell culture

All murine cell lines (pB2, pB3, EpCAM, Snail^hi^) and MCF7Ras human breast cancer cells containing doxycycline-inducible Luciferase or Slug and Sox9 constructs were a kind gift from Weinberg Lab and established and maintained as previously described (6, 21, 22, 55). Murine cell lines were cultured in a 1:1 mixture of Dulbecco’s modified Eagle’s medium (DMEM) and Ham’s F12 medium supplemented with 5% Bovine adult serum, 1x penicillin-streptomycin and 1x nonessential amino acids maintained at 37°C incubator containing 5% CO2. MCF7Ras cells were maintained in DMEM with 10% Bovine fetal serum and 1x penicillin-streptomycin as previously described (6, 21, 22, 55). All cell lines were routinely tested for mycoplasma (from 2019-2025) using the MycoAlert Mycoplasma Detection Kit (Lonza). All cell lines are negative for Mycoplasma and not authenticated since they were first acquired.

### Correlation analysis of EMP-related factors in canine tumors

Publicly-available datasets were accessed through GEO (61) or the Integrated Canine Data Commons (64, 65). Oral squamous cell carcinoma RNA-seq data was previously published (63). Genes that were significantly upregulated in two independent murine quasi-mesenchymal cell lines compared to epithelial counterparts and associated with immunomodulatory functions (22) were tested for anticorrelations with epithelial markers E-cadherin (*CDH1*) and EPCAM, EMT-TFs Snail and Slug (*SNAI1/2*), ZEB1/2, and TWIST1/2, and mesenchymal markers Vimentin (*VIM*) and N-cadherin (*CDH2*).

### Statistical analysis

All statistical analyses were performed using the GraphPad Prism v10.4.1 software. All count data represent standard error of the mean (SEM) using either two-tailed unpaired t-tests (Fig 4E, 4J) or an ordinary one-way ANOVA (Figs 1B-C, 2B-G, 3A-I, 4A, 4D, 4G, S2A-B, S4E-I). Survival analysis was performed using the Mantel-Cox Log Rank test (Fig 4F). Pearson correlation analysis was used to correlate normalized gene expression values (Figs 4B-C, 5, S5, S6). Asterisks indicate statistical significance where * = p<0.05, ** = p<0.01, *** p<0.001, **** = p<0.0001. Comparisons are not significant (p>0.05) if not indicated.

## Acknowledgments

We are grateful to all members of the Dongre Lab for their help in discussing and reviewing the manuscript. We thank Drs. Praveen Sethupathy, Jacquelyn Evans, Colleen Lau, and Robert Weiss for critical reading of the manuscript. We sincerely thank Dr. Jennifer Grenier and Faraz Ahmed from the Cornell University Biotechnology Resource Center Transcriptional Regulation and Expression Facility (RRID: SCR_022532) for performing the RNA sequencing. We are grateful to Dr. Yassi Hafezi of the Cornell University Biotechnology Resource Center Genomics Facility (RRID: SCR_021727) and Dr. Peter Schweitzer for their assistance in generating the scRNA-seq data. We thank all staff in the Cornell University Animal Health Diagnostic Laboratory for performing the immunohistochemical labeling of CMC samples. We thank Marco Hiller for assistance with the Aperio ScanScope Unit used to digitally scan slides. We thank Drs. Deborah Knapp and Deepika Dhawan for the access to canine invasive urothelial carcinoma RNA-seq data. Canine pulmonary carcinoma RNA-seq data was accessed through the National Cancer Institute Integrated Canine Data Commons. We thank Dr. Kelly Hume for helpful discussions regarding canine mammary carcinomas and reviewing the manuscript. Funding for this study was provided by the Judith Appleton PhD Early Career Excellence in Research Award (AD) and the Cornell University Center for Vertebrate Genomics Scholars Award (KMB).

## Author Contributions

Conceptualization: KMB, ADM, AD

Methodology: KMB, SN, LL, IN, IDV, SP, ADM, AD

Investigation: KMB, SN, LL, IN, IDV, SP, ADM, AD

Visualization: KMB

Supervision: ADM, SP, IDV, AD

Writing—original draft: KMB, AD

Writing—review & editing: KMB, SN, LL, IN, IDV, SP, ADM, AD

