## Supplementary Files for "Epithelial-Mesenchymal Plasticity is Associated with Immunosuppressive Features in Canine Mammary Carcinomas"

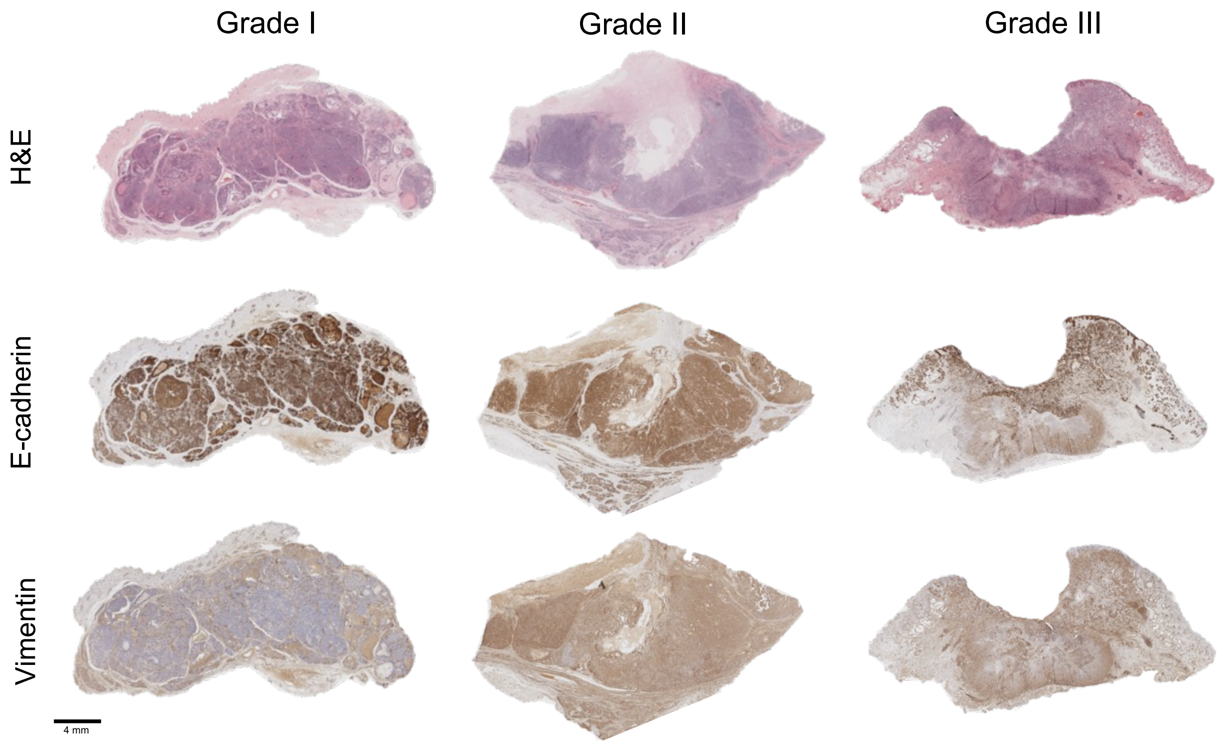


**Figure S1. Whole slide images of CMC labeled for EMP markers.** One CMC representative sample is provided for each grade. Samples were stained with hematoxylin and eosin (H&E) and labeled for EMP markers using E-cadherin and vimentin. Slides were then scanned using an Aperio CS2 ScanScope and the whole slide image is displayed. E-cadherin labeling is progressively lost, and vimentin co-expression increases with grade.

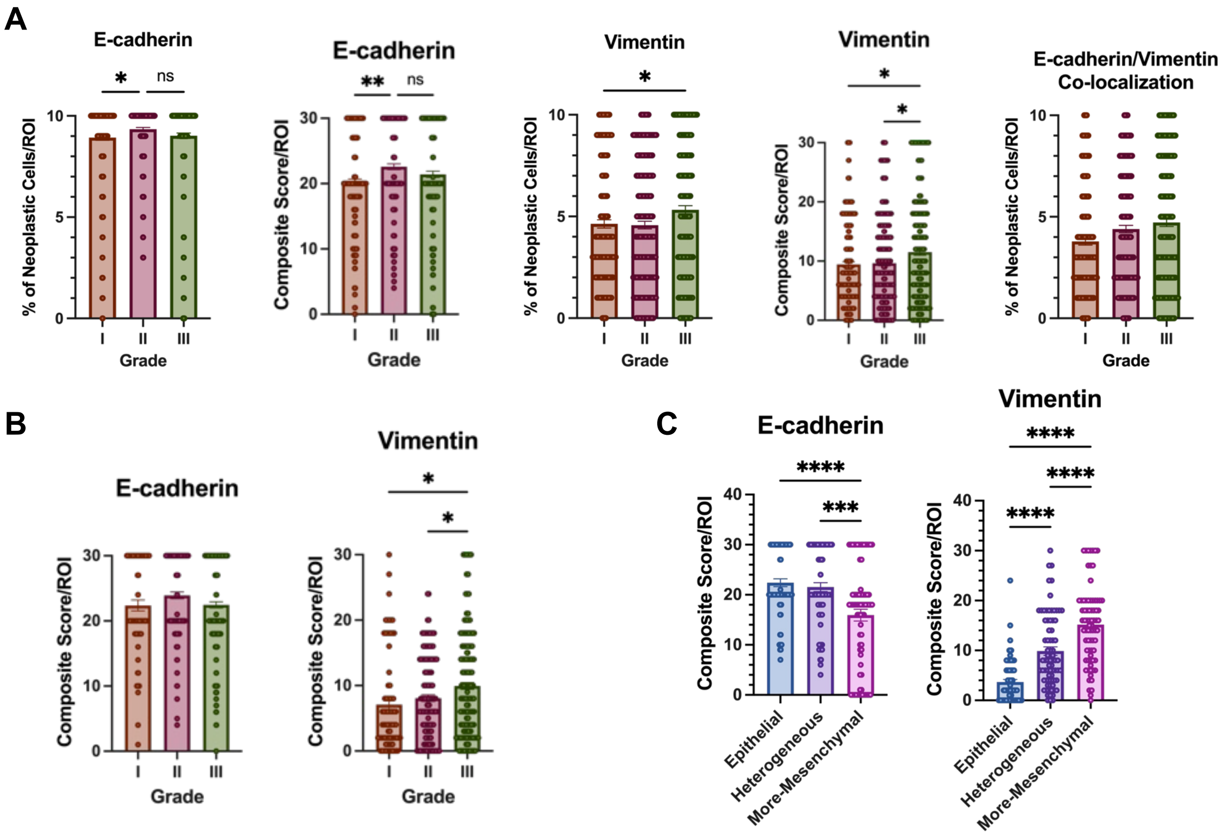


**Figure S2. Quantification of neoplastic cells expressing EMP markers in CMCs. (A)** Percentages of positive neoplastic cells for E-cadherin and vimentin labeling on a 0-10 scale, composite scores (product of % of positive cells and intensity of EMP marker labeling on a 0-3 scale) on a 0-30 scale, as well as estimated co-localization for all 52 samples. **(B)** Composite scores of E-cadherin and vimentin labeling in CMCs not displaying overt myoepithelial differentiation, on a 0-30 scale. **(C)** Composite scores of E-cadherin and vimentin labeling in CMCs selected for RNA-seq, on a 0-30 scale. All graphs are plotted as the mean value with error bars indicating the standard error of the mean. Data analyzed using one-way ANOVA **** $p < 0.0001$, *** $p < 0.001$, ** $p < 0.01$, * $p < 0.05$, ns ($p \geq 0.05$) if not indicated.

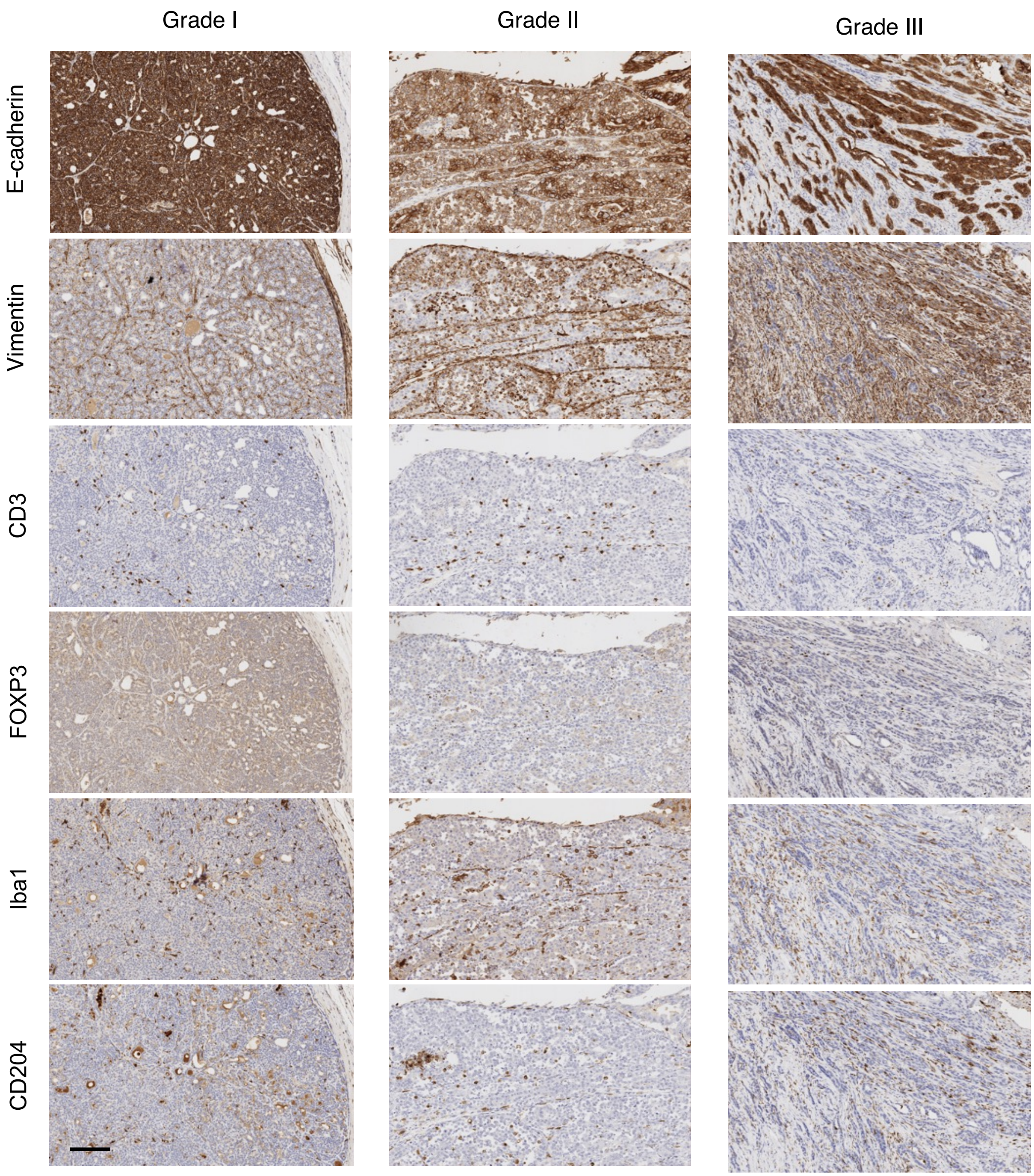


**Figure S3. Low magnification views of EMP and immune marker labeling of CMCs.** One CMC representative sample is provided for each grade, displaying the tumor center and periphery. Samples were labeled for EMP (E-cadherin, vimentin) and immune (CD3, FOXP3, Iba1, CD204) markers. Matched regions of interest are provided for each sample across all 6 markers. Scale bar indicates 200 μm.

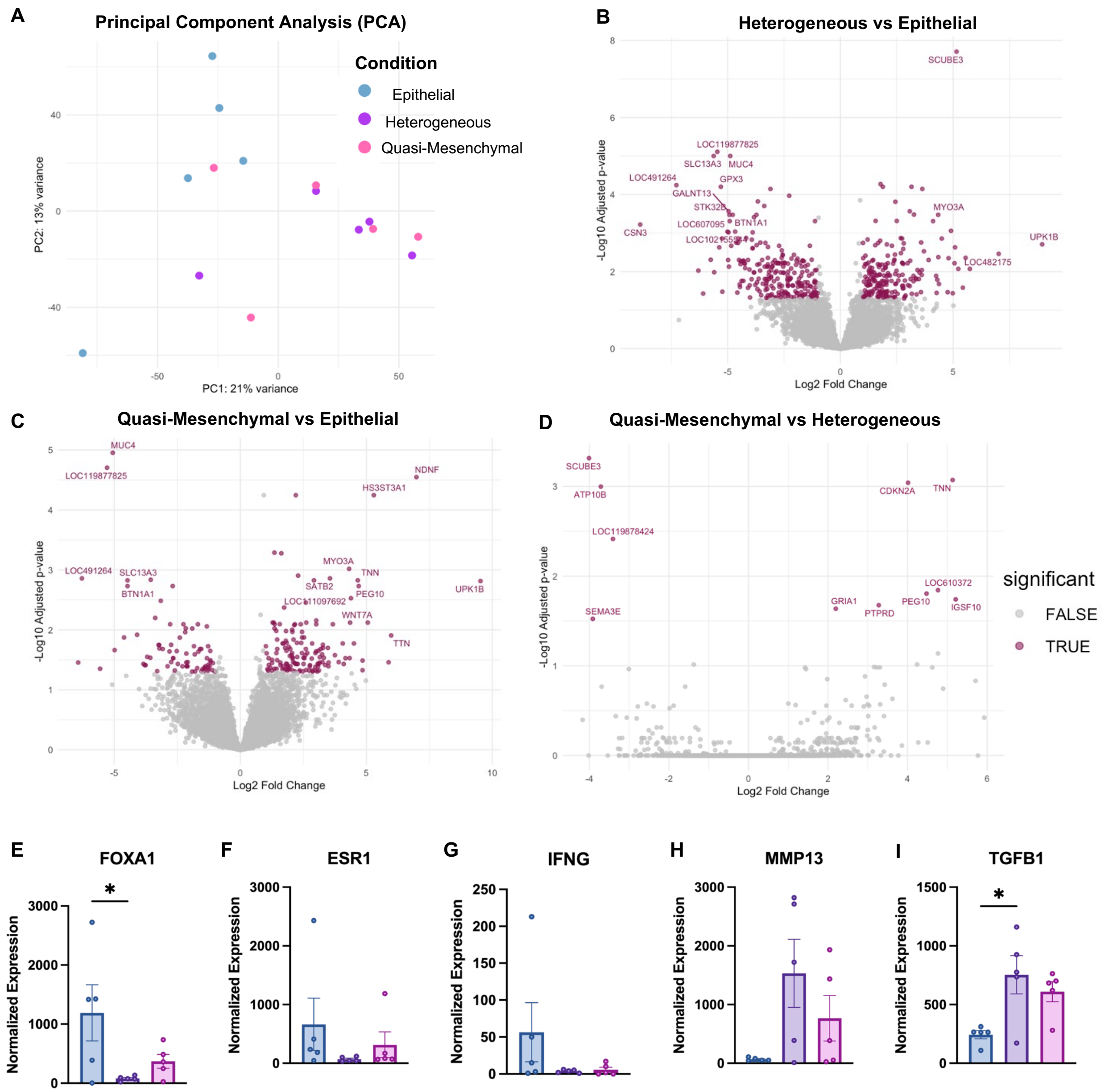


**Figure S4. RNA sequencing (RNA-seq) of epithelial, heterogeneous, and quasi-mesenchymal CMCs. (A)** Principal component analysis of samples used for RNA-seq. **(B-D)** Volcano plot of top 15 differentially expressed genes between epithelial, heterogeneous, and quasi-mesenchymal CMC samples. Differentially-expressed genes defined as genes with $|log_2FC| > 1$ and $p_{adj} < 0.05$. **(E-I)** Normalized expression of differentially-expressed genes of interest. Data analyzed using one-way ANOVA * $p < 0.05$, ns ($p \geq 0.05$) if not indicated.

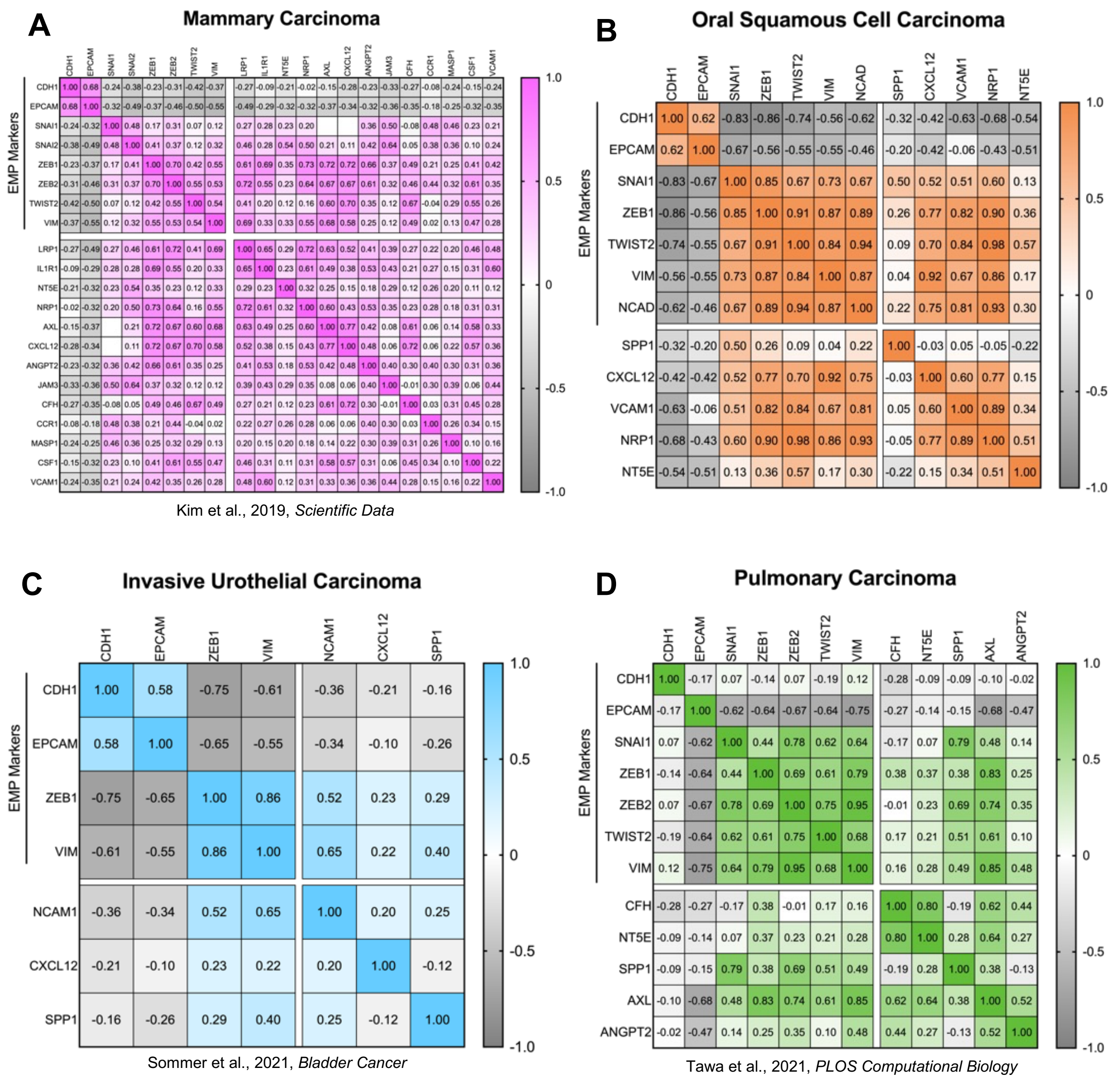


**Figure S5. Pearson correlation coefficients for EMP markers and immunosuppressive factors in canine carcinomas.** Correlation matrices of EMP markers and immunosuppressive factors that were upregulated in two independent MMTV-PyMT quasi-mesenchymal cell lines (pB3 and Snail$^{hi}$, Dongre et al., 2021, *Cancer Discovery,* Fig 5) across four canine cancers: **(A)** Mammary carcinoma, **(B)** Oral squamous cell carcinoma**, (C)** Invasive urothelial carcinoma, **(D)** Pulmonary carcinoma. Scale bars indicate Pearson r correlation values.

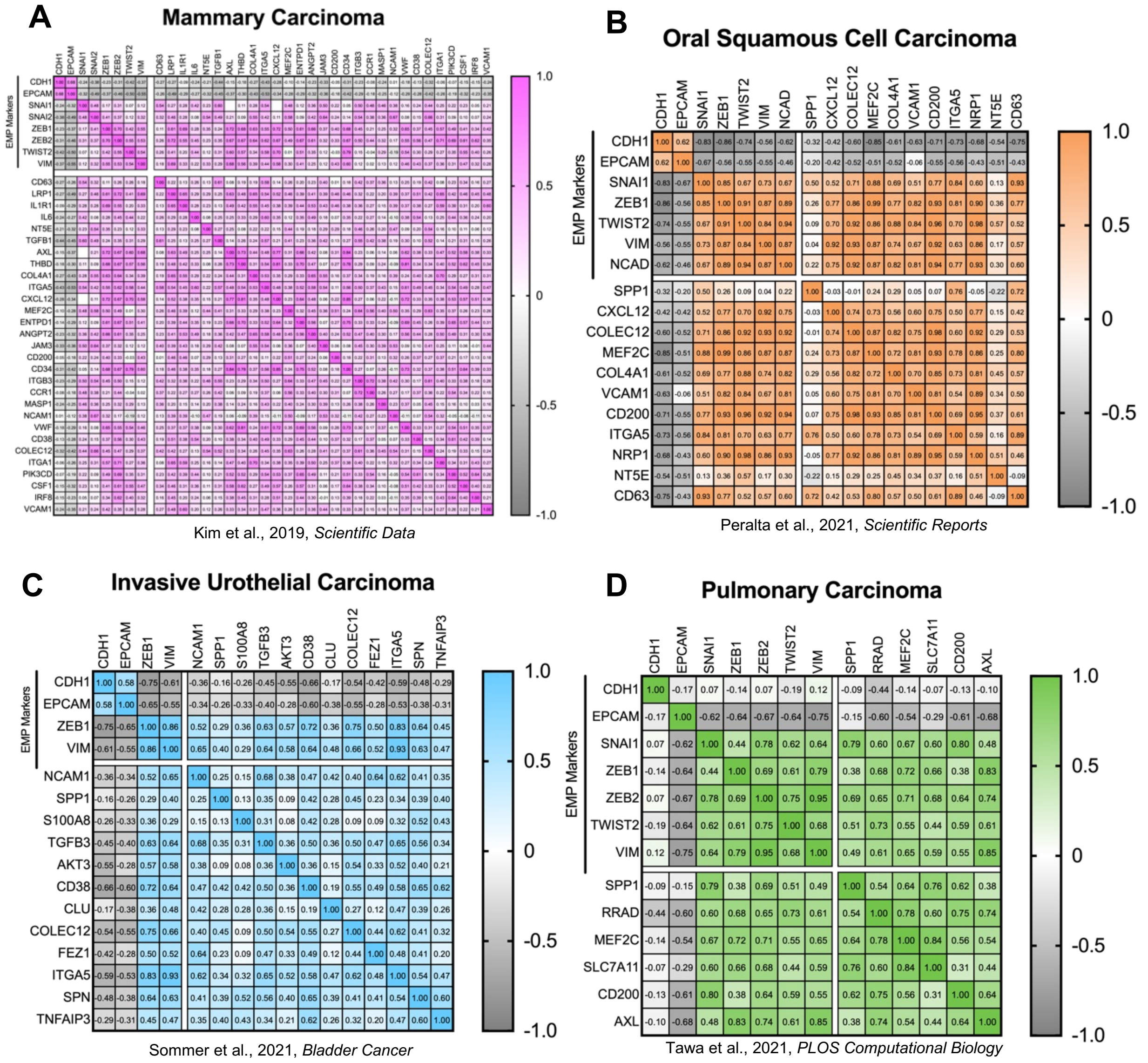


**Figure S6. Pearson correlation coefficients for EMP markers and extended list of immunosuppressive factors in canine tumors.** Correlation matrices of EMP markers and immunosuppressive factors that were upregulated in either of two independent MMTV-PyMT quasi-mesenchymal cell lines (pB3 and Snail$^{hi}$, Dongre et al., 2021, *Cancer Discovery*) across four canine cancers: **(A)** Mammary carcinoma, **(B)** Oral squamous cell carcinoma, **(C)** Invasive urothelial carcinoma**,** **(D)** Pulmonary carcinoma. Scale bars indicate Pearson r correlation values.

**Table S1. Canine mammary carcinoma sample information.** NOS = Not otherwise specified. F = Female, FS = Female spayed.

| Sample number | Age (years) | Sex | Breed | Diagnosis | Vascular invasion (Y/N) | Grade | Selected for RNA-seq | RNA-seq condition |
|---|---|---|---|---|---|---|---|---|
| **1** | 6 | FS | NOS | Complex carcinoma | N | 1 | Yes | Epithelial |
| **2** | 6 | FS | NOS | Complex carcinoma | N | 1 | | |
| **3** | 3 | FS | Mixed breed | Mixed carcinoma | N | 1 | | |
| **4** | 5 | F | Irish Wolfhound | Tubulopapillary carcinoma | N | 1 | Yes | Epithelial |
| **5** | 12 | FS | Yorkshire terrier | Complex carcinoma | N | 1 | | |
| **6** | 12 | F | Mixed breed | Complex carcinoma | N | 1 | | |
| **7** | 8 | FS | Yorkshire terrier | Tubulopapillary carcinoma | N | 1 | | |
| **8** | 6 | F | Pitbull terrier | Intraductal solid cribriform carcinoma | N | 1 | Yes | Epithelial |
| **9** | 8 | FS | Mixed breed | Solid and tubular carcinoma | N | 1 | Yes | Heterogeneous |
| **10** | 10 | FS | Mixed breed | Mixed carcinoma | N | 1 | Yes | Quasi-Mesenchymal |
| **11** | 9 | FS | Mixed breed | Tubulopapillary carcinoma | N | 1 | Yes | Heterogeneous |
| **12** | 6 | FS | Yorkshire terrier | Tubular carcinoma | N | 1 | Yes | Quasi-Mesenchymal |
| **13** | 3 | FS | Terrier, NOS | Tubulopapillary carcinoma | Y | 1 | | |
| **14** | 4 | FS | Maltese | Complex carcinoma with papillomatosis | N | 1 | | |
| **15** | 9 | F | Mixed breed | Mixed carcinoma | N | 2 | | |
| **16** | 10 | FS | Boston Terrier | Intraductal papillary carcinoma | N | 2 | | |
| **17** | 11 | FS | Field Spaniel | Solid carcinoma | N | 2 | | |
| **18** | 10 | FS | Rhodesian Ridgeback | Intraductal papillary carcinoma | N | 2 | | |
| **19** | 10 | FS | Miniature Schnauzer | Complex carcinoma | N | 2 | | |
| **20** | 15 | F | Maltese | Solid and tubular carcinoma | N | 2 | | |
| **21** | 15 | F | Maltese | Mixed carcinoma | Y | 2 | | |
| **22** | 3 | FS | Mixed breed | Intraductal solid carcinoma | N | 2 | | |
| **23** | 11 | F | Chihuahua | Tubulopapillary carcinoma | Nodal metastasis | 2 | | |
| **24** | 10 | FS | Mixed breed | Complex carcinoma | N | 2 | | |
| **25** | 11 | F | Golden Retriever | Tubulopapillary carcinoma | Nodal metastasis | 2 | Yes | Epithelial |
| **26** | 6 | FS | Pitbull terrier | Tubulopapillary carcinoma | Y | 2 | | |
| **27** | 11 | FS | American Staffordshire Terrier | Tubular carcinoma | N | 2 | | |
| **28** | 9 | FS | Maremma sheepdog | Solid carcinoma | N | 2 | | |
| **29** | 12 | FS | Mixed breed | Solid carcinoma | Y | 2 | | |
| **30** | 11 | FS | Yorkshire terrier | Mixed carcinoma | Y | 2 | | |
| **31** | 4 | FS | American Staffordshire Terrier | Intraductal solid carcinoma with squamous differentiation | Y | 2 | | |
| **32** | 11 | F | Chihuahua | Mixed carcinoma | N | 2 | | |
| **33** | 10 | F | Maltese | Tubular carcinoma | N | 3 | | |
| **34** | 13 | FS | Mixed breed | Solid and tubular carcinoma | N | 3 | | |
| **35** | 8 | FS | Mixed breed | Inflammatory carcinoma | Y; also nodal metastasis | 3 | Yes | Heterogeneous |
| **36** | 11 | FS | Mixed breed | Anaplastic carcinoma | Y | 3 | | |
| **37** | 8 | F | Saluki | Simple tubular carcinoma | N | 3 | | |
| **38** | 8 | FS | German Shepherd | Solid anaplastic carcinoma | Y | 3 | Yes | Quasi-Mesenchymal |
| **39** | 12 | FS | Mixed breed | Malignant myoepithelioma with comedonecrosis | Y | 3 | | |
| **40** | 10 | FS | Yorkshire terrier | Anaplastic carcinoma with osteosarcomatous differentiation | N | 3 | | |
| **41** | 12 | FS | Miniature Pinscher | Solid carcinoma with comedonecrosis | Y | 3 | | |
| **42** | 8 | F | Goldendoodle | Anaplastic carcinoma with squamous differentiation | Y | 3 | | |
| **43** | 12 | FS | Sled dog | Complex carcinoma | N | 3 | | |
| **44** | 8 | FS | Mixed breed | Intraductal tubulopapillary carcinoma | Y | 3 | Yes | Epithelial |
| **45** | 9 | FS | Mixed breed | Tubular carcinoma | Y | 3 | Yes | Heterogeneous |
| **46** | 12 | FS | Mixed breed | Tubular carcinoma | Y | 3 | Yes | Heterogeneous |
| **47** | 11 | F | NOS | Mixed carcinoma with osteosarcomatous differentiation | Y | 3 | Yes | Quasi-Mesenchymal |
| **48** | 11 | FS | Dachshund | Complex carcinoma with comedonecrosis | N | 3 | | |
| **49** | 5 | FS | NOS | Solid and tubular carcinoma | N | 3 | | |
| **50** | 9 | FS | German Shepherd | Complex carcinoma | N | 3 | | |
| **51** | 8 | F | Beagle hound | Solid carcinoma | Y | 3 | | |
| **52** | 9 | FS | Mixed breed | Tubular carcinoma | N | 3 | Yes | Quasi-Mesenchymal |
